# Extracting allelic read counts from 250,000 human sequencing runs in Sequence Read Archive

**DOI:** 10.1101/386441

**Authors:** Brian Tsui, Michelle Dow, Dylan Skola, Hannah Carter

## Abstract

The Sequence Read Archive (SRA) contains over one million publicly available sequencing runs from various studies using a variety of sequencing library strategies. These data inherently contain information about underlying genomic sequence variants which we exploit to extract allelic read counts on an unprecedented scale. We reprocessed over 250,000 human sequencing runs (>1000 TB data worth of raw sequence data) into a single unified dataset of allelic read counts for nearly 300,000 variants of biomedical relevance curated by NCBI dbSNP, where germline variants were detected in a median of 912 sequencing runs, and somatic variants were detected in a median of 4,876 sequencing runs, suggesting that this dataset facilitates identification of sequencing runs that harbor variants of interest. Allelic read counts obtained using a targeted alignment were very similar to read counts obtained from whole genome alignment. Analyzing allelic read count data for matched DNA and RNA samples from tumors, we find that RNA-seq can also recover variants identified by WXS, suggesting that reprocessed allelic read counts can support variant detection across different library strategies in SRA. This study provides a rich database of known human variants across SRA samples that can support future meta-analyses of human sequence variation.

## 1. Introduction

The reduction of sequencing cost in recent years^1^ has allowed researchers to progress from sequencing and analyzing a single reference human genome to studying the individual genomes of thousands of subjects^2^. The large number of sequencing studies being conducted, together with journal publication requirements for authors to deposit raw sequencing runs in a centralized and open access sequencing archive like Sequence Read Archive (SRA)^3^ have made it possible to perform large scale data analysis on the millions of publically-available sequencing runs.

The SRA contains raw sequencing runs from a variety of projects from large scale consortium studies including Epigenome Roadmap^4^, ENCODE^5^, The 1000 Genomes Project^2^, to small studies being conducted by various independent laboratories. However, the publicly available raw sequencing data are large in size which translates into high storage and computational requirements that hinder access for the broader research community. These requirements can be somewhat mitigated by using preprocessed data such as gene expression matrices, ChIP-seq peak files, or summarized variant information, as such files are much smaller in size. For example, the 1000 Genomes project, The Cancer Genome Atlas (TCGA)^6^ and Genotype-Tissue Expression project (GTEx)^7^ all offer summarized variant information extracted from the raw sequences in Variant Call Format (VCF) files, containing allelic read counts for both reference and alternative alleles and base quality information which could be used for variant calling.

There have been many efforts to reprocess raw sequencing reads to a more tractable form. However, many of the SRA data reprocessing efforts^8,9^ have focused on quantifying gene expression using public RNA-seq data deposited in the SRA. Sequencing data also capture information about sequence variants, raising the possibility of studying patterns of genetic variation using the SRA. The possibility of extracting variants from RNA-seq was demonstrated on a small scale in a 2015 study^10^ where the authors extracted variants using the GATK RNA-seq variant calling pipeline on 5,499 RNA-seq runs in the SRA.

Variant calling typically requires multiple user-specified parameters such as a minimum cut-off for total or read-specific coverage, and usually attempts to model sequencing error explicitly. The primary information used in variant detection is the allelic fraction, the proportion of sequencing reads that support the variant position. Read mapping is highly concordant between alignment tools like bowtie^11^, bwa^12^, novoalign^13^, supporting the idea, at least for DNA and RNA sequencing experiments, estimates of allelic fraction should be fairly consistent regardless of the specific alignment tool. Using a conservative set of known genetic variants that are unlikely to be the result of sequencing errors, simple filters on coverage or allelic fraction should be sufficient to control error rates at acceptable levels. This would make it possible to collect and analyze known variants across the SRA without applying more complex variant callers.

To explore this possibility, we constructed an allelic read count extraction pipeline to systematically reprocess all available sequencing runs from the SRA. We first applied standard quality filtering to the unaligned reads (see Methods) and then aligned the reads to a subset of the human reference genome that covers 390,000 selected somatic and germline variants curated by the NCBI dbSNP^14^ using bowtie2^11^. To show that this targeted reference does not introduce unwanted biases into the alignment step, we validated our pipeline performance against alignments performed using whole reference genomes. We next used the TCGA sample-matched Whole Exome Sequencing (WXS) and RNA-seq cohort to confirm that allelic read counts derived from RNA-seq accurately recover variants detected by WXS. We then applied this pipeline to systematically extract variants from over 250,000 sequencing runs in the SRA. Finally, we demonstrated that this allelic read count resource can be used to investigate variants in RNA sequencing studies, even at the single cell level.

## 2. Results

### 2.1. Building a fast allelic fraction extraction pipeline for the SRA

As of the end of 2017 the SRA included data from 10,642 human sequencing studies consisting of 697,366 publicly available sequencing runs, encompassing various library strategies such as RNA-seq, whole exome sequencing (WXS), whole genome sequencing (WGS), and ChIP-seq (Methods) and this number continues to increase at a rapid pace (Fig. 1). All of the human sequences deposited in the database were derived from samples carrying germline and somatic variants from the corresponding biospecimen regardless of the original study designs. This presents the opportunity to perform meta-analysis of human genetic variation across studies in the SRA.

**Fig. 1.**
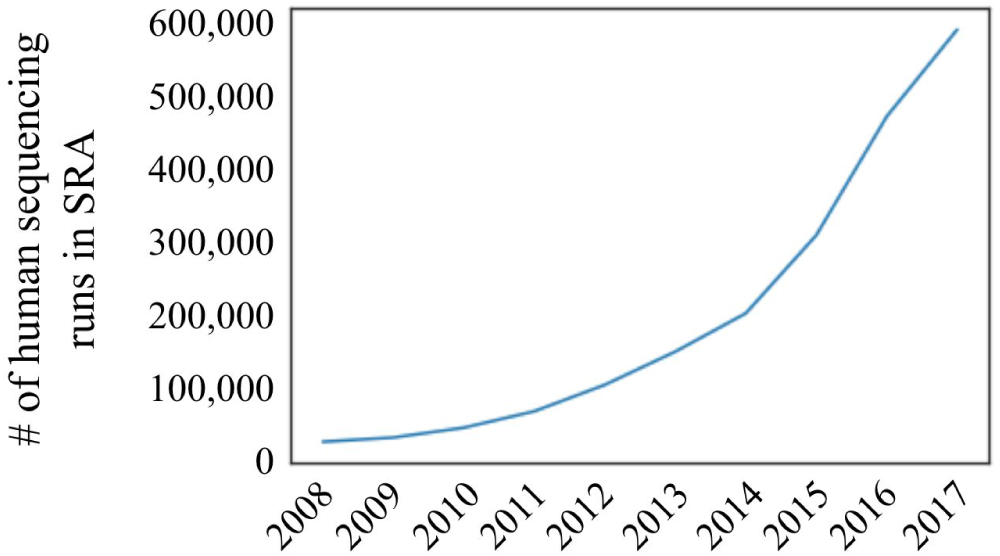
Number of human sequencing runs are increasing exponentially in the SRA

However, the complete SRA spans over 1,835 trillion bases, introducing both computational and storage resource requirements that would hinder most researchers from conducting a meta-analysis across many sequencing studies. Therefore, to enable efficient secondary analysis for researchers with limited access to high performance computing (HPC) infrastructure, we sought to process this vast amount of data into a form that can fit on a 1 TB hard disk. To accomplish this, we developed an efficient data processing pipeline (Fig. 2).

**Fig. 2.**
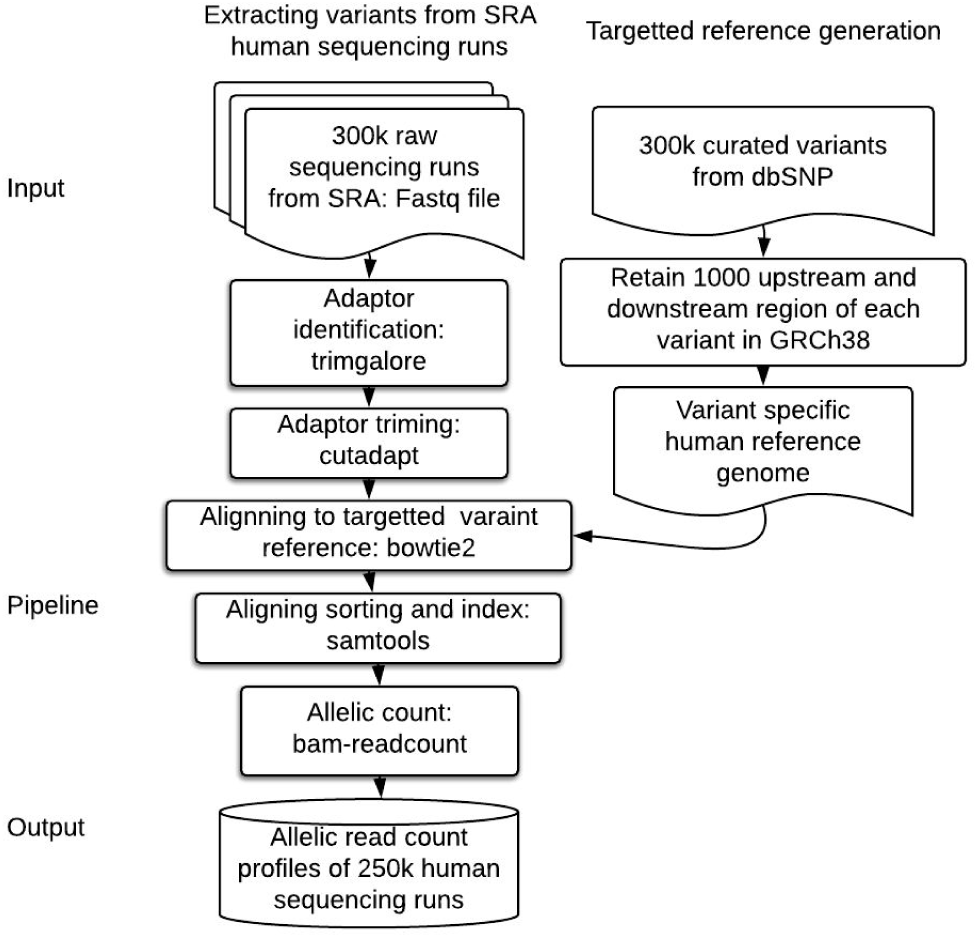
Simple pipeline for extracting human sequencing runs from SRA

We first created a targeted alignment reference that focuses on regions that harbor known variants (n=393,242) curated by NCBI dbSNP^14^. These consist predominantly of variants with PubMed references or that have been referenced in selected variant databases (OMIM, LSDB, TPA, or in NCBI curated as diagnostic related). The variants consist mostly of missense mutations with synonymous and truncating mutations accounting for about 15% of the database. Most are germline variants, although the dataset includes a small set of curated somatic mutations^15^. The characteristics of the variants are summarized in Table 1.

**Table 1.** Key characteristics of variants in targeted reference

| Property | Variant type | Number of variants | % of variants |
| --- | --- | --- | --- |
|  | All | 393,242.00 |  |
|  | Has 3D structure. SNP3D table | 20,800.00 | 5.29 |
| Resource link property | Cited by PMC article | 170,292.00 | 43.30 |
|  | Cited in PubMed or referenced in a clinical database | 201,900.00 | 51.34 |
| Substitution type | Non-synonymous missense | 91,827.00 | 23.35 |
|  | synonymous | 32,778.00 | 8.34 |
|  | Non-synonymous frameshift | 17,824.00 | 4.53 |
|  | Nonsense mutation | 9,286.00 | 2.36 |
| Genotype properties | Genotypes available, also on high density Genotyping kit and have phenotype associations present in dbGaP | 148,114.00 | 37.66 |
| Phenotype properties | Submitted from a locus-specific database | 141,029.00 | 35.86 |
|  | Has OMIM/OMIA | 59,617.00 | 15.16 |
|  | Somatic (not germline) variant | 37,704.00 | 9.59 |

We created the reference alignment index by masking the reference to exclude DNA sequences outside of a region spanning the 1000 base pairs upstream and 1000 base pairs downstream of each variant. This filtering method had been first adopted by Deng *et al*. to optimize sequencing data processing turnaround times^16^.

### 2.2. Large scale allelic read count extraction of human sequence data

We retained only sequencing runs from the top five library strategies (RNA-seq, WGS, WXS, AMPLICON, ChIP-seq), and sequencing runs with more than 150 million bases sequenced (equivalent to at least three million reads if the samples have 50 bp per read), corresponding to a total of 304,939 sequencing runs. Of these, 253,005 were successfully processed (Fig. 3) without error with 300 cpu-cores in 30 days. Library strategies were divided between paired-end (64.8%) and single-end (35.2%) sequencing. The difference between the number of pair-end sequencing and single-end sequencing reflects the differing needs of various experimental designs (Table 2). For example, paired-end sequencing greatly improves the identification of splice isoforms in RNA-seq and structural variants in exome-seq, whereas it provides fewer benefits for other library types that would justify the increased cost relative to single-end sequencing.

**Table 2.**
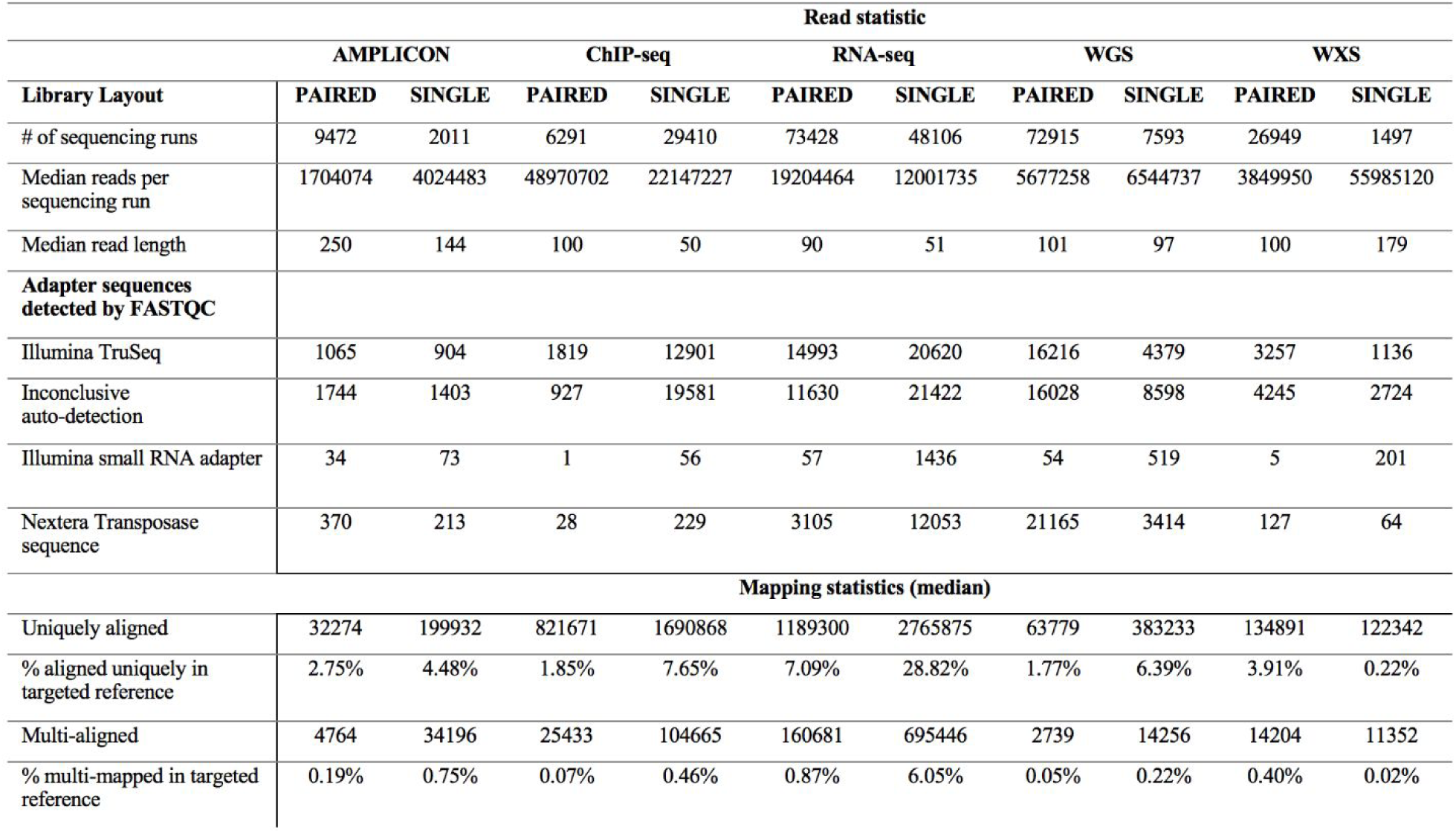
Characteristics and mapping statistics of the SRA sequencing runs

| Library Layout | Read statistic |  |  |  |  |  |  |  |  |  |
| --- | --- | --- | --- | --- | --- | --- | --- | --- | --- | --- |
|  | AMPLICON |  | ChIP-seq |  | RNA-seq |  | WGS |  | WXS |  |
|  | PAIRED | SINGLE | PAIRED | SINGLE | PAIRED | SINGLE | PAIRED | SINGLE | PAIRED | SINGLE |
| # of sequencing runs | 9472 | 2011 | 6291 | 29410 | 73428 | 48106 | 72915 | 7593 | 26949 | 1497 |
| Median reads per sequencing run | 1704074 | 4024483 | 48970702 | 22147227 | 19204464 | 12001735 | 5677258 | 6544737 | 3849950 | 55985120 |
| Median read length | 250 | 144 | 100 | 50 | 90 | 51 | 101 | 97 | 100 | 179 |
| **Adapter sequences detected by FASTQC** |  |  |  |  |  |  |  |  |  |  |
| Illumina TruSeq | 1065 | 904 | 1819 | 12901 | 14993 | 20620 | 16216 | 4379 | 3257 | 1136 |
| Inconclusive auto-detection | 1744 | 1403 | 927 | 19581 | 11630 | 21422 | 16028 | 8598 | 4245 | 2724 |
| Illumina small RNA adapter | 34 | 73 | 1 | 56 | 57 | 1436 | 54 | 519 | 5 | 201 |
| Nextera Transposase sequence | 370 | 213 | 28 | 229 | 3105 | 12053 | 21165 | 3414 | 127 | 64 |
| **Mapping statistics (median)** |  |  |  |  |  |  |  |  |  |  |
| Uniquely aligned | 32274 | 199932 | 821671 | 1690868 | 1189300 | 2765875 | 63779 | 383233 | 134891 | 122342 |
| % aligned uniquely in targeted reference | 2.75% | 4.48% | 1.85% | 7.65% | 7.09% | 28.82% | 1.77% | 6.39% | 3.91% | 0.22% |
| Multi-aligned | 4764 | 34196 | 25433 | 104665 | 160681 | 695446 | 2739 | 14256 | 14204 | 11352 |
| % multi-mapped in targeted reference | 0.19% | 0.75% | 0.07% | 0.46% | 0.87% | 6.05% | 0.05% | 0.22% | 0.40% | 0.02% |

**Fig. 3.**
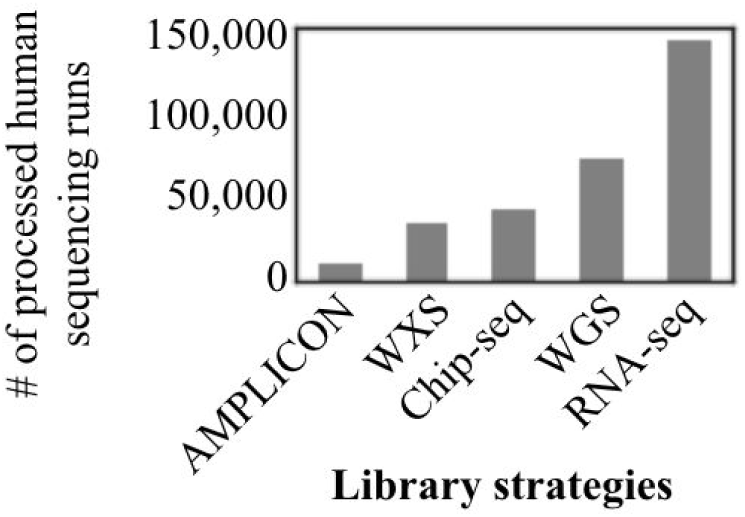
Distribution of processed SRA data

One utility that emerges from reprocessing the sequencing data is for imputing experimental annotations. For example, the SRA metadata is not standardized to contain important experimental variables like read length or adaptor sequences, however this information can be easily determined from the raw sequences. A median read length of 95 bp was observed. Most runs (206,360 = 81.56%) had adaptors automatically detected and removed. Sequence and mapping statistics are detailed in Table 2. Over these sequencing runs, a median of 2.98% of base pairs were identified as adaptors and were removed. A median base quality Phred score of 36 was observed, suggesting a high overall quality of the sequenced bases in the SRA.

Overall, a median of 296.3 million bases and 10,044,529 read fragments per sample were observed. A median of 5.83% of the reads were aligned to the targeted variant regions (**Methods**). Adding read length, adaptor contents, number of reads and percentage aligned to the metadata allows the user to better understand the quality of the sequencing runs and filter them accordingly.

### 2.3. Pipeline performance for targeted variant detection

To assess the accuracy of allelic read counts extracted from this targeted reference we compared counts obtained through our pipeline to those extracted from samples pre-aligned to the complete hg38 genome index and downloaded directly from the TCGA. We also took advantage of matched DNA/RNA sequencing in TCGA to evaluate the extent to which allelic read counts extracted from RNA-seq reflect the variants detected from WXS (See section 2.5). We used 524 whole exome tumor sequences from the TCGA Low Grade Glioma (LGG) dataset to assess the performance of our pipeline, as this dataset included the well-known variant (IDH1 R132H) which could serve as a positive control.

The reads from each tumor were aligned to the targeted SNP index and the allelic read counts were compared to the pre-generated alignments available from the TCGA. The resulting variant-locus-by-nucleotide read count matrix contains the read count for each of the four nucleotides across the 393,242 targeted variants at 387,950 genomic sites. We then flattened the nucleic base read count matrix into a single allelic read count vector. For each sample, we compared allelic read counts for all variants obtained using alignment to a targeted reference against allelic read counts obtained from the existing TCGA alignments to a complete reference. Read counts were highly correlated. Figure 4A shows an example from a single TCGA tumor (UUID: 2b0048e0-a062-40d2-a1e1-4bb763ea0ead), in which a median of 98.2% variants differed less than one log_2_ fold change in allelic read count from the existing alignment (95% confidence interval: 0.0088 - 0.0554). We found similar correlation across all 524 samples, with a median Pearson correlation (R) of 0.98 for the allelic read counts (95% CI: 0.928 - 0.992; Fig. 4B).

**Fig. 4.**
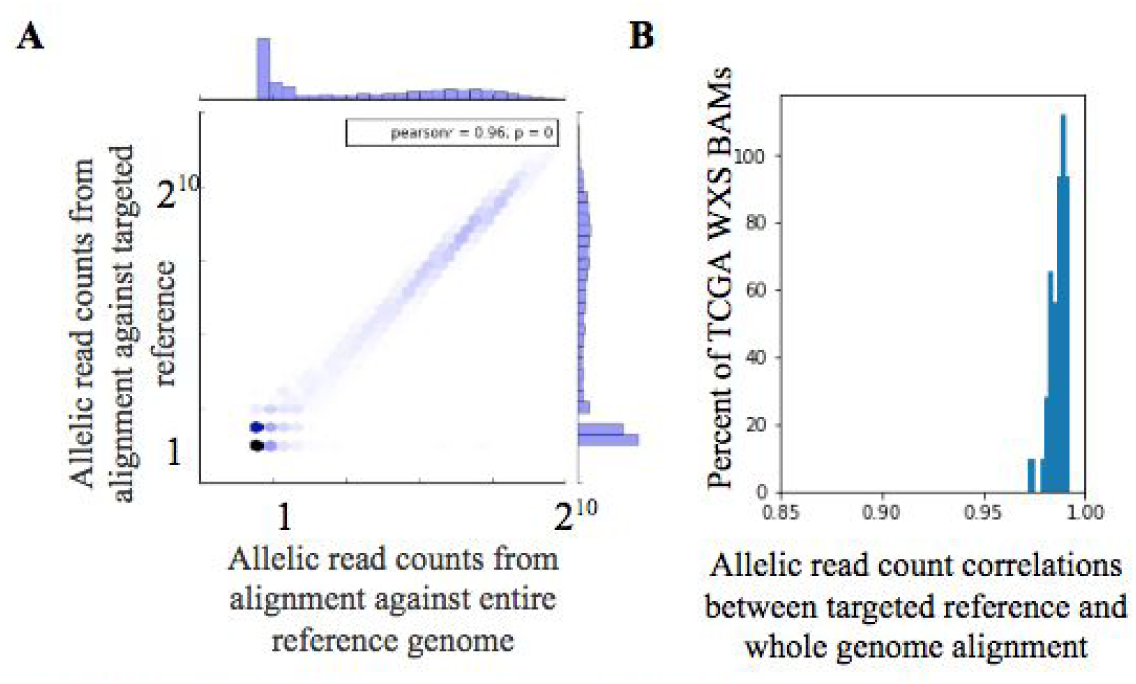
Targeted reference remains accurate for sequence alignment. **A** Allelic read count correlation between whole genome aligned. **B** Distribution of allelic read count correlations.

### 2.4. Effects of PCR duplicates on estimating allelic fraction

We next evaluated the necessity of removing putative PCR duplicate reads after alignment based on the extent to which such duplicates bias the estimate of allelic fraction in TCGA. Although most sequence alignment pipelines include a step for removing duplicate reads that result from PCR amplification, recent studies have cast doubt on the benefit of doing so for variant analysis^17,18^. Also, naively removing the duplicated reads could result in overcorrection in high coverage sequencing^19^.

We therefore investigated the effect of sequence duplicate removal for all 300k targeted variants across the 524 samples. We compared the allelic read counts extracted with and without duplicate removal for each tumor WXS alignment, and observed a median correlation of 0.983 (95% CI: 0.983-0.990), suggesting duplicate removal had limited impact on allelic read counts. However, we did observe a substantial bias in allelic read count estimates when duplicates are included among sites with very high sequence read coverage. Figure 5A shows an example using UUID: 0e2c395e-ddda-4833-b1ee-31a9bd08a845. In this sample, deduplicated allelic read counts recover 88.9% of the original allelic read counts among all the variants with ≤100 reads support, while the deduplicated allelic only recover 33.7% of the original allelic read count among all the variants with >100 reads, a 2.63 fold reduction in read count extracted from in the high coverage region (Fig. 5A, slope of grey bar and red bar respectively). Nonetheless, across all 524 samples we observed a difference in allelic fraction < 0.05 for over 90% of the variants when duplicates were excluded, except in extreme cases with over 10,000 mapped reads (median 0.4% of the variants) (Fig. 5B). Thus with high quality sequencing data, filtering duplicates should result in only minor improvement to the data.

**Fig. 5.**
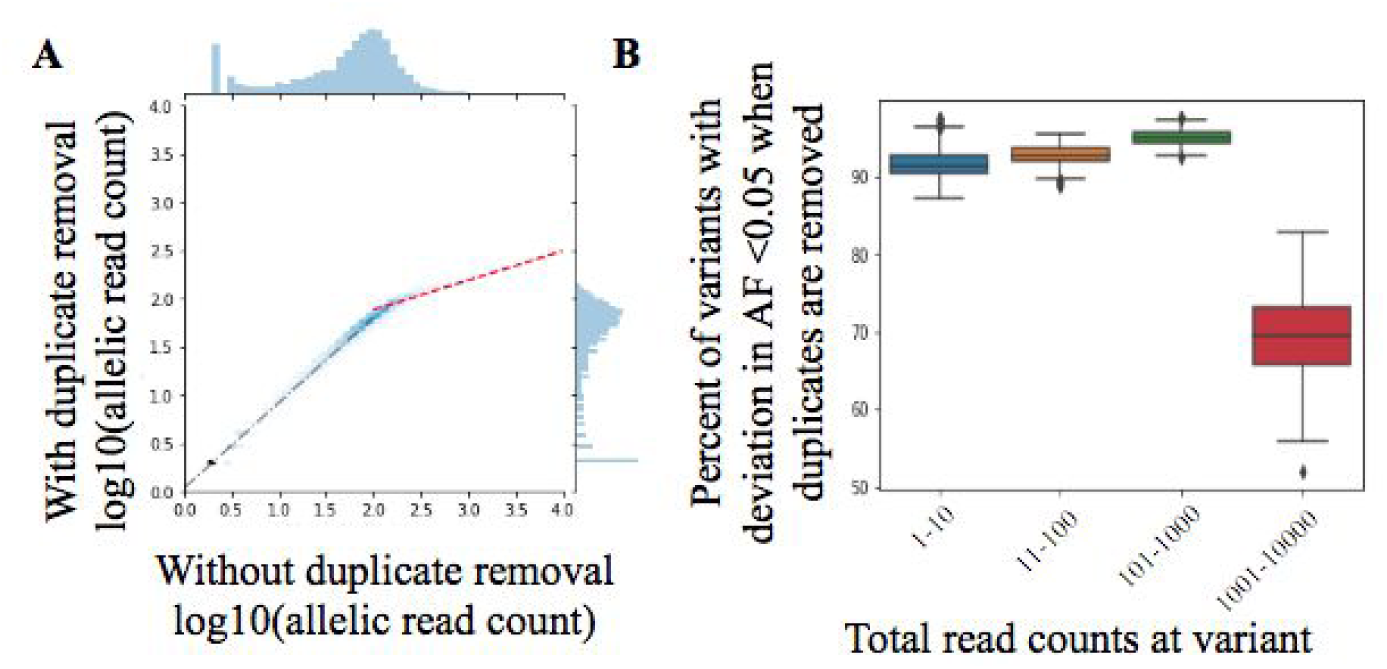
**A** Duplicate removal doesn’t scale in high read depth region. **B** allelic traction are comparable regardless of duplicate removal except in sites with extremely high read count.

### 2.5. Evaluating variant extraction from RNA-seq using matched DNA/RNA samples

The SRA includes over 100k RNA-seq runs and these data contain information about the variant status of the transcribed DNA. To determine the extent to which variants can be extracted from RNA-seq by our pipeline, we first compared allelic fractions between matched exome sequencing on the one hand and RNA sequencing data in TCGA on the other. TCGA contains samples which have been subjected to both WXS and RNA-seq, which makes it a natural resource for comparing the performance of variant calls derived from RNA-seq data using the WXS-derived variants. We evaluated the possibility of using allelic read counts from RNA-seq to detect both germline and somatic variants.

To evaluate the reliability of allelic read counts for identifying germline variants in RNA sequence reads, we first compared read fractions for germline variants that were homozygous in the corresponding TCGA WXS sample. After collecting all sites that had at least 10 reads and were homozygous for the variant allele in the WXS read data, we evaluated the read counts at those same sites in the RNA-seq data. A median of 5827 sites had at least 10 reads to support the variant in both WXS and RNA-seq for each sample. Across all samples, a median of 97% (95% CI: 95.5% - 97.9%) of sites that were homozygous in the DNA were also found to be homozygous in the matched RNA-seq data.

Next, we explored the utility of allelic read counts for identifying somatic mutations from RNA sequencing data. First, as a positive control, we evaluated the hotspot IDH1 somatic mutation on chromosome 2:208248388 with 395G>A in the template strand, which is most prevalent somatic variant in TCGA LGG on WXS as called by Varscan ^20^ (n=371, 70.80% of patients). This variant had been previously identified as enriched in LGG tumors and its status is a major molecular prognostic factor in glioma as noted by the World Health Organization (WHO)^21^. Using the 524 LGG tumors, we estimated allelic composition using read counts in the matched RNA-seq and WXS independently with our pipeline. The IDH1 mutation status in WXS exhibits a bimodal distribution (Fig. 6A). We selected 10 reads as the cutoff for defining a positive WXS variant. The reference allele was detected in the WXS in all tumors, and 351 patients also had the alternative allele. Over these patients the RNA-seq achieved an area under the precision recall curve (AUPRC) of 0.98 in detecting IDH1 variants observed in the WXS data (Fig. 6B).

**Fig. 6.**
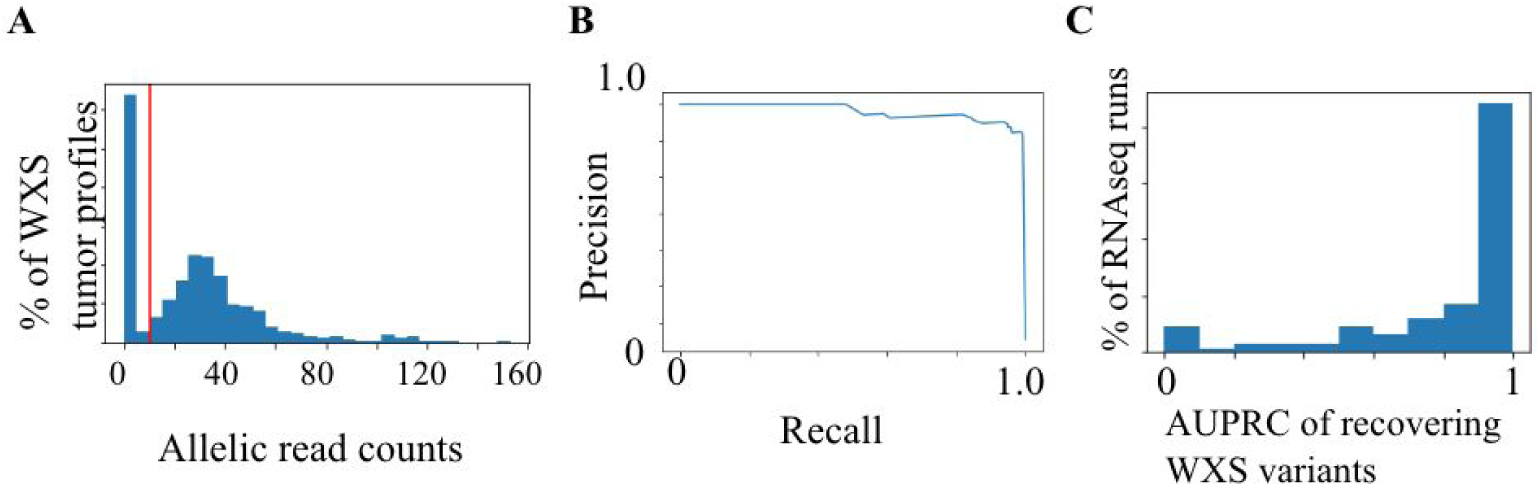
RNA-seq can recover variants extracted from WXS. **A** Minor allelic read counts of 1DH1 hotspot mutation. Vertical red line is the binomial distribution cutoff (10 read counts). **B** distribution of minor allele of IDH1 (395C>T in template strand). **C** RNAseq has high area under the precision recall curve (AUPRC) of recovering WXS variants

We next evaluated the top 100 most frequently observed somatic variants reported by TCGA in the LGG samples that also coincided with the targeted variants, since recurrent mutations are more likely to be drivers and present the most attractive therapeutic targets^22^. We used the Precision Recall Curve (PRC) framework to determine the extent to which allelic read counts supported expression of the mutant allele. RNA-seq generally recapitulated WXS variants (Fig. 6C), with 70% of the variants having an AUPRC > 0.8, suggesting that majority of the variants called by exome sequencing are expressed in the tumor. However, we do observe 6% of the variants with an AUPRC less than 0.1 when their presence was predicted from RNA-seq allelic fraction. Importantly, these later variants were found in fewer than 10 WXS samples, such that the most recurrent somatic mutations are also more frequently consistently expressed. Thus while absence of a somatic variant cannot be definitively determined from RNA-seq (mutations can be present but not expressed), the most recurrent variants appear to be frequently expressed, suggesting that many somatic mutations of interest will be detectable in RNA-seq data from cancer studies deposited in the SRA.

### 2.6. Variant landscape of the SRA

After validating the general reliability of our allelic fraction estimates, we analyzed 300K variants across the SRA. Properties of the variants are listed in Table 1. Of 300K variants, 170,292 were referenced by PubMed and 138,559 were curated by NCBI as clinically-relevant variants. Out of 156,757 variants with annotated functional effects, the majority were missense mutations (n=91,827). Also, 37,704 variants were annotated as somatic mutations, derived from cancer studies. Overall, the data included a median of three variants per gene across 21,889 genes.

We collected read counts for reference and alternative alleles at these 300K positions for 253,005 human sequencing samples in the SRA. We used default minimum threshold of two reads^23^ as the cut-off for Varscan^20^. The distribution of the number of variants are shown in Figure 7. Known germline variants were detected in a median of 912 sequencing runs, known somatic variants were detected in a median of 4,876 sequencing runs, and known reference alleles were detected in a median of 33,232 sequencing runs. 337 somatic variants, 3,068 germline variants and 23,044 reference alleles were covered by at least two reads in more than half of the sequencing runs, suggesting that SRA data can be repurposed for studying many variants. To facilitate the analysis of variants, we collected allelic read count in each SRA sample into a table (see Data Availability). This read count file allows researchers to rapidly identify which sequencing runs in the SRA have read support for a particular variant.

**Fig. 7.**
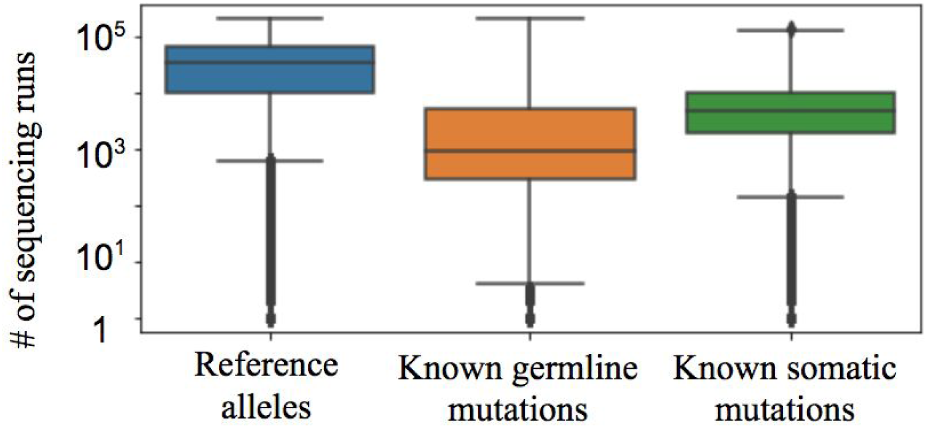
Distribution of variants detected associated with each variant type

### 2.7. Extracting unannotated single cell variants in cancer in SRA

Genotype annotations are often missing or incomplete in the SRA, and this limits the reusability of the SRA data. Here, we show that, using the reprocessed data, we were able to recover an important oncogenic mutation BRAF V600E in a single cell RNA-seq study of a patient with myeloid leukemia at diagnosis and as well as at three and six months after diagnosis ^24^.

Traditional variant calling relies on high sequencing depths to provide the statistical power to make confident calls. However, since each cell carries only two copies of each chromosome, the low recovery of single cell sequencing makes variant calling from DNA resequencing difficult. Since RNA also contains information about underlying variants and may exist at hundreds of copies per cell^25^, calling variants from single-cell RNA-seq data may circumvent the limitations of DNA resequencing for variants in transcribed regions.

We were able to detect an important oncogenic mutation, BRAF V600E, in single cells using our unified allelic read counts. The overall read depth for the region was 45.9 reads and 17 sites within the 20 bp windows around BRAF V600E had read support for the reference allele. Alternative alleles at the BRAF V600 hotspot were detected in more than 95% cells (Fig. 8A). Also, the alternative allele (T) had a median base quality Phred score of 38 (Fig. 8B) and a median of 22.0 reads to support it (Fig. 8C). Interestingly, we observed a reduction in the reference allele read count over the course of treatment (Fig. 8D) with a corresponding higher fraction of reads supporting the alternate allele, suggesting that the clone with BRAF mutations became more prevalent among the surviving cancer cells, concording with the observation in the study that relapse occurred after treatment.

**Fig. 8.**
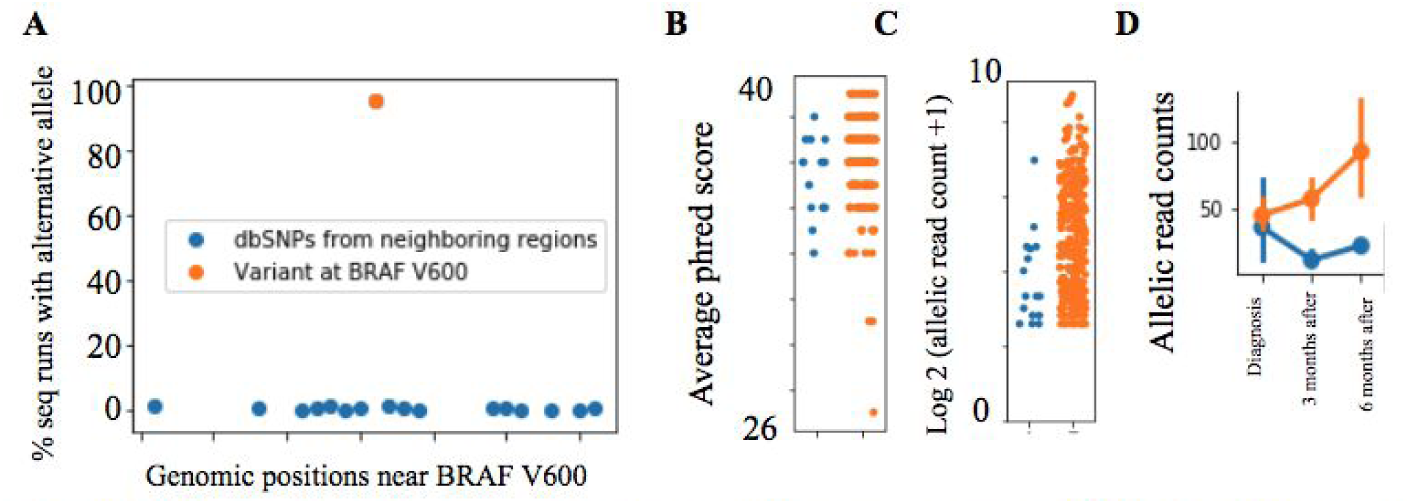
**A** allelic read counts can recover obvious variants (example: chr7-140753336). **B** Base quality, and **C** read count of reads at chr7-140753336 for reference allele (blue) and alternative allele (orange). **D** Allelic read count of alternative allele can track cancer progression.

## 3. DISCUSSION

Most published studies on non-protected raw sequencing data are expected to be deposited in the NCBI SRA as a result of journal requirements, and this vast amount of raw sequencing data represents a an opportunity to power large-scale meta-analyses for the interaction of sequence variants with experimental conditions. However, these petabytes worth of sequencing data introduce a computational challenge for analyzing such variants. One solution is to develop a map of relevant sequence variants in the SRA using allelic count profiles.

To create allelic read count profiles from the SRA, we constructed a bioinformatics pipeline with minimal processing turnaround time by mapping the raw sequencing reads to a targeted reference specific to key somatic and germline variant(s) curated by the NCBI dbSNP. We validated the accuracy of the pipeline by comparing read counts obtained with targeted alignment to counts obtained using complete alignment pipelines, and evaluated genotype consistency across multiple sequencing datasets derived from the same sample. These results confirm that the targeted alignment pipeline generates allelic read counts that are highly correlated to those from whole genome alignments.

Variant calling has traditionally been performed from DNA sequences, but WXS and WGS library strategies comprise only 40% of the total human SRA data. Thus we also sought to infer the presence of variants from RNA-seq allelic read counts. While RNA may be less reliable for inferring the presence or absence of variants due to gene and allele-specific expression, 61.8% of the RNA-seq samples have more than a million reads mapped onto the targeted variant regions. We also found that highly recurrent somatic mutations detected in WXS of low grade gliomas were also frequently expressed in matched RNA-seq data.

To the best of our knowledge, this is the first attempt to massively reprocess the human samples in the SRA for the purpose of extracting allelic read counts. The computational infrastructure required to generate variant data at scale presents a barrier to many researchers. Consortia that generate a large volume of sequencing data, such as GTEX, TCGA or the 1000 Genome Project, all offer preprocessed files that enable researchers from the broader community to identify novel findings. Although variant calls are available for some of the datasets included in SRA, significant effort would be required to aggregate these disparate datasets, and most of the non-consortia SRA samples do not have such data available. Simply providing allelic read counts derived through a common bioinformatic pipeline also avoids technical variation that can result from different choice of computational tools and their associated parameter choices. Therefore, we contend that our unfiltered allelic read counts will have broad utility for *post hoc* analysis.

Many applications require estimates of the magnitude of allelic fraction for inference. This would be particularly useful for questions related to imprinting or reconstruction of tumor subclonal architecture. We found that presence of duplicate reads did not significantly bias estimates of allelic fraction when the quality of the sequencing data is high. However for lower quality datasets or different library strategies, it may still be necessary to remove duplicate reads to obtain high quality estimates. Further analysis is merited to determine which datasets or variants are most confounded if duplicates are not removed. Future releases of the database will include estimates of allelic fraction both before and after removing PCR duplicates.

In conclusion, by reprocessing the raw sequencing runs from the SRA, we improve the findability, accessibility, interoperability, reusability (FAIR) ^26^ of of 250,000 sequencing runs. As the SRA continues to grow, it will be necessary to continuously update the map of variants present in SRA samples. To support variant meta-analyses using the SRA, the next requirement will be unification of the SRA data, including biospecimen and experimental annotations. We anticipate that further refinement of the SRA through efforts such as this will promote reanalysis of existing datasets and lead to new biological discoveries.

## 4. METHODS

### 4.1. SRA Metadata download

SRA metadata (files: NCBI_SRA_Metadata_Full_.tar.gz and SRA_Run_Members.tab) were downloaded from ftp.ncbi.nlm.nih.gov/sra/reports/Metadata/ on Jan 4 2018. These files contain the raw freetext biospecimen and experimental annotations. SRA_Run_Members.tab details the relationships between SRA project ID (SRP), sample ID (SRS), experiment ID (SRX) and sequencing run IDs (SRR). We processed only sequencing runs with accession visibility status “public”, with availability status “live”, and sequencing runs that contains more than 150 million nucleotides bases. We also only included sequencing runs generated from the following library strategies: RNA-Seq, WGS, WXS, ChIP-Seq, AMPLICON. Only samples with layout defined as either SINGLE or PAIRED were considered. We removed SRA study ERP013950 as we noticed it has annotation indicating a total of 85,608 WGS sequencing runs which seem to stem from erroneous submission, as it was only associated with nine biological samples (BioSample) IDs and the experimental annotation was unclear on the nature of the study.

### 4.2. NCBI dbSNP structure

NCBI dbSNP uses a bitfield encoding schema with each bit in the byte indicating support by a type of evidence (ftp://ftp.ncbi.nlm.nih.gov/snp/specs/dbSNP_BitField_latest.pdf). Some evidence types are derived from databases, for example, Online Mendelian Inheritance in Man (OMIM, url: https://www.omim.org/), Locus-Specific DataBases (LSDB, url: http://www.hgvs.org/locus-specific-mutation-databases), and Third Party Annotation (TPA, url: https://www.ddbj.nig.ac.jp/ddbj/tpa-e.html). OMIM contains the genotypes and phenotypes of all known mendelian disorders for over 15,000 human genes. LSDB provides gene-centric links to various databases that collect information about variant phenotypes. TPA is a nucleotide sequence data collection assembled from experimentally determined variants from DDBJ, EMBL-Bank (https://www.ebi.ac.uk/), GenBank, International Nucleotide Sequence Database Collaboration (INSDC) (http://www.insdc.org/), and/ or Trace Archive (https://trace.ncbi.nlm.nih.gov/Traces/home/) with additional feature annotations supported by peer-reviewed experimental or inferential methods.

### 4.3. Targeted reference building

Variants were obtained from dbSNP (downloaded on 4, January on 2017 from ftp://ftp.ncbi.nlm.nih.gov/snp/organisms/human_9606_b150_GRCh38p7/VCF/00-All.vcf.gz), which contained 325,174,853 sites in total, effectively one tenth of our selected human reference genome length (3,099,734,149 bp, version: hg38). We retained only variants with a resource link to any of the existing databases or with support from NCBI curation, indicated by a non zero value for byte 2 of Flag 1 in the NCBI bit field encoding schema, resulting in 393,242 variants. To generate a targeted reference for these variants, we defined 1000 bp downstream and 1000 bp upstream of each SNP as the mapping window. All the regions outside of the windows were masked with base “N” using bedtools v2.26.0 in the reference FASTA file. The reference index was built using bowtie2 v2.2.6^11^ with the merged FASTA file, using default parameters.

### 4.4. Extracting variants from raw sequencing read FASTQ file

We used SRA^3^ prefetch v2.8.0 to download SRR files. Next, fastq-dump v2.4.2 from SRA tool kit was used to extract FASTQ files from SRR into the standard output stream. Trim Galore! version 0.4.0 (url: https://github.com/FelixKrueger/TrimGalore) was then applied to identify adapter sequences using the first 10,000 reads, and the identified adaptor sequence was trimmed in the FASTQ file using cutadapt version 1.16^27^, the trimmed reads were then aligned onto the targeted reference (we did not use Trim Galore! to trim the adaptor as it cannot be easily UNIX piped). Bowtie2 was run with the “--no-unal” parameter to retain only the reads mappable to the target regions in order to minimize the amount of aligned reads for sorting. The alignment file was than sorted using samtools v1.2. and samtools idxstats was used for calculating the number of reads that mapped onto each FASTA reference record. bam-readcount v0.8.0 was used for extracting the per-base allelic read count and per-base quality in the sorted alignment file for each of the targeted genomic coordinates. The paired-end reads were processed the same way as the single-end reads with the exception that paired-end and interleave reads options in fastq-dump, cutadapt, and bowtie2, were specified to ensure proper treatment of paired-end reads.

### 4.5. TCGA download

A gdc_manifest was downloaded from the gdc portal on 2017-12-27. We downloaded the TCGA data using gdc-client v1.3.0. We downloaded the associated metadata using the TCGA REST API interface https://api.gdc.cancer.gov/files/. All the alignment files preprocessed from TCGA using GATK pipeline were downloaded. The alignment files were mapped onto GRCh38 with all the raw reads, including read sequence duplicates.

## 5. Code and data availability

The python scripts for the pipeline and the jupyter-notebooks for generating the figures are deposited on github (https://github.com/brianyiktaktsui/Skymap) and the data is publicly available on synapse (https://www.synapse.org/#!Synapse:syn11415602).

## 6. Acknowledgments

We thank all members of the Carter, Mesirov and Ideker lab for scientific feedback and comments. The results here are partly based upon the data generated by the TCGA Research Network: http://cancergenome.nih.gov/. This work was funded by NIH grants DP5-OD017937, RO1 CA220009 and a CIFAR fellowship to H.C. Preprint of this article is submitted for consideration in Pacific Symposium on Biocomputing © 2018 copyright World Scientific Publishing Company.

## References

1. Wetterstrand, K. A. DNA sequencing costs: data from the NHGRI Genome Sequencing Program (GSP). (2013).

2. The 1000 Genomes Project Consortium. A global reference for human genetic variation. Nature 526, 68 (2015).

3. Leinonen, R., Sugawara, H., Shumway, M. & International Nucleotide Sequence Database Collaboration. The sequence read archive. Nucleic Acids Res. 39, D19–21 (2011).

4. Roadmap Epigenomics Consortium et al. Integrative analysis of 111 reference human epigenomes. Nature 518, 317–330 (2015).

5. ENCODE Project Consortium. An integrated encyclopedia of DNA elements in the human genome. Nature 489, 57–74 (2012).

6. Cancer Genome Atlas Research Network et al. The Cancer Genome Atlas Pan-Cancer analysis project. Nat. Genet. 45, 1113–1120 (2013).

7. Carithers, L. J. & Moore, H. M. The Genotype-Tissue Expression (GTEx) Project. Biopreserv. Biobank. 13, 307–308 (2015).

8. Collado-Torres, L. et al. Reproducible RNA-seq analysis using recount2. Nat. Biotechnol. 35, 319–321 (2017).

9. Lachmann, A. et al. Massive Mining of Publicly Available RNA-seq Data from Human and Mouse. bioRxiv 189092 (2017). doi:10.1101/189092

10. Deelen, P. et al. Calling genotypes from public RNA-sequencing data enables identification of genetic variants that affect gene-expression levels. Genome Med. 7, 30 (2015).

11. Langmead, B. & Salzberg, S. L. Fast gapped-read alignment with Bowtie 2. Nat. Methods 9, 357–359 (2012).

12. Li, H. & Durbin, R. Fast and accurate short read alignment with Burrows-Wheeler transform. Bioinformatics 25, 1754–1760 (2009).

13. Ruffalo, M., LaFramboise, T. & Koyutürk, M. Comparative analysis of algorithms for next-generation sequencing read alignment. Bioinformatics 27, 2790–2796 (2011).

14. Kitts, A. & Sherry, S. The Single Nucleotide Polymorphism Database (dbSNP) of Nucleotide Sequence Variation. (National Center for Biotechnology Information (US), 2011).

15. Classes of Genetic Variation Included in dbSNP. (National Center for Biotechnology Information (US), 2005).

16. Deng, J. et al. Targeted bisulfite sequencing reveals changes in DNA methylation associated with nuclear reprogramming. Nat. Biotechnol. 27, 353–360 (2009).

17. Ebbert, M. T. W. et al. Evaluating the necessity of PCR duplicate removal from next-generation sequencing data and a comparison of approaches. BMC Bioinformatics 17 Suppl 7, 239 (2016).

18. Stratford, J. et al. Abstract 5276: Impact of duplicate removal on low frequency NGS somatic variant calling. Cancer Res. 76, 5276–5276 (2016).

19. Zhou, W. et al. Bias from removing read duplication in ultra-deep sequencing experiments. Bioinformatics 30, 1073–1080 (2014).

20. Koboldt, D. C. et al. VarScan 2: somatic mutation and copy number alteration discovery in cancer by exome sequencing. Genome Res. 22, 568–576 (2012).

21. Louis, D. N. et al. The 2016 World Health Organization Classification of Tumors of the Central Nervous System: a summary. Acta Neuropathol. 131, 803–820 (2016).

22. Chang, M. T. et al. Identifying recurrent mutations in cancer reveals widespread lineage diversity and mutational specificity. Nat. Biotechnol. 34, 155–163 (2016).

23. Xu, C. A review of somatic single nucleotide variant calling algorithms for next-generation sequencing data. Comput. Struct. Biotechnol. J. 16, 15–24 (2018).

24. Giustacchini, A. et al. Single-cell transcriptomics uncovers distinct molecular signatures of stem cells in chronic myeloid leukemia. Nat. Med. 23, 692–702 (2017).

25. Albayrak, C. et al. Digital Quantification of Proteins and mRNA in Single Mammalian Cells. Mol. Cell 61, 914–924 (2016).

26. Wilkinson, M. D. et al. The FAIR Guiding Principles for scientific data management and stewardship. Scientific Data 3, 160018 (2016).

27. Martin, M. Cutadapt removes adapter sequences from high-throughput sequencing reads. EMBnet.journal 17, 10–12 (2011).

